# MurineCyto-Det: A High-Resolution Murine BALF Cytology Dataset for Leukocyte Segmentation and Detection

**DOI:** 10.64898/2026.05.08.723893

**Authors:** Thang X. Le, Lan-Anh T. Tran, Dia A. Farabi, Shiyuan Wang, Anh T.Q. Phan, Stephania A. Cormier, Anvi Taada, Derek McGrew, Yuxuan Du, Luan D. Vu

**Affiliations:** Department of Civil Engineering, University of Minho, Guimarã es, 4800-058, Portugal; Department of Mathematics, Computer Science and Statistics, Ghent University, Gent, 9000, Belgium; Department of Electrical Engineering, The University of Texas at San Antonio, San Antonio, Texas, 78249, USA; Pennington Biomedical Research Center, Louisiana State University, Louisiana, 610101, USA; Department of Molecular and Microbiology Immunology, The University of Texas at San Antonio, Texas, 78249, USA

## Abstract

Automated analysis of murine bronchoalveolar lavage fluid (BALF) cytology is important for preclinical respiratory research, yet progress has been limited by the lack of publicly available, well-annotated mouse BALF image datasets. We present MurineCyto-Det, a high-resolution murine BALF cytology dataset comprising 333 image tiles of size 1024×1024 pixels, annotated across five cytological categories with both pixel-level segmentation masks and one-to-one matched bounding boxes. The dataset contains 14,551 annotated cell instances and supports two complementary analysis tasks: morphology-oriented cell segmentation and object-level cell detection. To establish reproducible benchmark baselines, we evaluated representative segmentation and detection models. The results demonstrate the practical utility of MurineCyto-Det while highlighting realistic challenges arising from class imbalance, small object size, irregular cell morphology, and ambiguous debris-like structures. MurineCyto-Det provides a standardized resource for developing, evaluating, and comparing automated methods for murine BALF cytology analysis. The dataset is publicly available at https://doi.org/10.5281/zenodo.17608677.

## 1 Introduction

Bronchoalveolar lavage fluid (BALF) provides a direct and informative window into pulmonary physiology and pathology. In both clinical and experimental settings, BALF cytological analysis remains indispensable to assess inflammation, infection, and injury in the lower respiratory tract^1–4^. In mouse models-central to immunology, toxicology, and respiratory disease research-BALF analysis serves as a primary readout for lung injury and immune cell recruitment^5–9^. Changes in specific cell populations often mirror characteristic pathophysiological states: lymphocytosis in hypersensitivity pneumonitis or sarcoidosis^10,11^, neutrophilia in infection or acute respiratory distress^12,13^, and eosinophilia in allergic inflammation. Consequently, quantitative and morphological analyzes of BALF cells are critical to interpret disease mechanisms and therapeutic outcomes.

Traditionally, BALF cytology has relied on cytospin preparations followed by manual microscopic examination, where trained personnel identify and count monocytes, macrophages, lymphocytes, neutrophils and eosinophils^5,14^. Although considered a gold standard, this process is labor-intensive, and difficult to scale. This highlights the need for an automated method to support less experienced individuals in improving efficiency when working with BALF cells. Recent advances in artificial intelligence (AI), particularly deep learning, have transformed morphological diagnostics in human cytology^15–18^. Convolutional neural networks (CNNs) now achieve expert-level performance in automated recognition of hematological and cytological cells, including bone marrow smears^17^, peripheral blood^19,20^ and human BALF samples^18,21,22^. However, despite their potential, comparable resources for murine BALF, one of the most widely used tools in respiratory research, are conspicuously absent. Current AI models and datasets focus almost exclusively on human clinical samples, leaving a significant gap for preclinical and translational studies. Without standardized, publicly available murine BALF datasets and validated benchmarks, progress toward reproducible automated cytological analysis in basic research remains limited.

To address this unmet need, we present MurineCyto-Det, the high-resolution, pixel-level annotated dataset dedicated to automated detection and classification of murine BALF cells.The data set comprises images acquired under controlled experimental conditions, encompassing the main types of leukocytes: monocytes/macrophages, lymphocytes, neutrophils and eosinophils along with additional debris and background annotations to reflect realistic cytospin variability. Each image includes both bounding-box and segmentation-mask annotations, enabling dual benchmarking for object detection and morphological segmentation. Together with standardized training, validation, and test splits, MurineCyto-Det establishes a common platform for quantitatively consistent and reproducible benchmarking of algorithmic performance in automated murine BALF cytology.

In addition to releasing the dataset, we conducted benchmark evaluations to demonstrate the utility of MurineCyto-Det for automated BALF cytology analysis. We focused on two complementary tasks: pixel-level segmentation and object-level detection. Segmentation enables detailed assessment of cell boundaries, morphology, and spatial structure, supporting morphology-based quantification and phenotype characterization. Detection provides a more efficient framework for localizing, classifying, and counting individual cells, making it suitable for scalable cytology screening workflows. Together, these tasks capture the major analytical needs of automated BALF cytology and provide standardized reference points for future method evaluation. By introducing MurineCyto-Det, we aim to support reproducible AI-based analysis of murine BALF cytology and facilitate the development of robust computational tools for preclinical respiratory research.

## 2 Related work

### 2.1 Existing cytology datasets and benchmark gaps

High-quality annotated image datasets are essential for developing and objectively benchmarking automated cytology analysis methods. Existing public resources have primarily focused on human peripheral blood or human clinical BALF cytology. For peripheral blood analysis, the BCCD dataset^23^ provides bounding-box annotations for cell detection, while more detailed datasets such as Raabin-WBC^24,25^ and LISC^26^ include annotations that support leukocyte segmentation and classification. Other public hematology datasets^27,28^ have also contributed to algorithm development for blood-cell analysis. However, these resources differ substantially in imaging conditions, annotation protocols, class definitions, and acquisition settings, which limits direct comparison across studies and restricts their transferability to specialized cytology settings such as murine BALF^29^. BALF cytology datasets are more limited and have mainly been developed from human clinical samples. Recent human BALF resources have provided valuable high-resolution images and annotations for AI-based BALF cell analysis^18^. Nevertheless, human BALF datasets do not fully capture the morphology, staining characteristics, cell-density patterns, and cytospin artifacts observed in murine samples. These differences create a domain gap between human and mouse BALF cytology and limit the direct use of human-trained models in preclinical mouse studies. Despite the importance of mouse models in respiratory research, no publicly available dataset has been dedicated specifically to murine BALF cytology with both pixel-level segmentation masks and matched object-level bounding boxes across major leukocyte categories. This absence has limited reproducible algorithm development and standardized performance comparison for automated murine BALF cytology. MurineCyto-Det addresses this gap by providing high-resolution murine BALF cytology images with expert-validated annotations and paired segmentation and detection labels.

### 2.2 Motivation and contribution

Automated BALF cytology requires both morphology-oriented and object-level analysis. Pixel-level segmentation provides detailed information about cell boundaries, shape, and local morphology, which is important for morphometric analysis and phenotype characterization. Object detection, in contrast, localizes and classifies individual cells with bounding boxes, making it more suitable for efficient cell counting, screening, and differential cell analysis. A murine BALF dataset that supports both tasks is therefore needed to reflect the complementary analytical requirements of automated cytology workflows. To address this need, we introduce MurineCyto-Det, a publicly available murine BALF cytology dataset containing high-resolution images, expert-validated annotations across five biologically relevant cell categories, pixel-level segmentation masks, and corresponding bounding boxes. The paired annotation design enables both detailed segmentation-based analysis and efficient detection-based analysis on the same image set. In parallel with the dataset release, we provide representative benchmark baselines for two core computer vision tasks: pixel-level segmentation and object-level detection. These baselines are intended to demonstrate the practical use of MurineCyto-Det and to provide standardized reference points for future method evaluation and comparison, rather than to serve as an exhaustive ranking of available architectures. By combining expert-validated murine BALF annotations with standardized benchmark tasks, MurineCyto-Det provides a reproducible foundation for automated cytology analysis in preclinical respiratory research.

## 3 Dataset construction

### 3.1 Data acquisition

BALF samples were collected from 5–6-week-old BALB/c mice according to the Guide for the Care and Use of Laboratory Animals. The protocol was approved by the Institutional Animal Care and Use Committee at Louisiana State University and Pennington Biomedical Research Center, and the University of Texas at San Antonio (UTSA), all Association for Accreditation of Laboratory Animal Care accredited.

Mice were intranasally exposed to either respiratory syncytial virus (RSV) at 2 ×10^5^ TCID_50_ per gram body weight (TCID_50_ : 50% tissue culture infectious dose), influenza A/PR8/34 (H1N1) at 1 × 10^4^ plaque-forming units (PFU) per mouse, or sterile saline as a control. At 6 days post-infection or exposure, BALF was collected via tracheal intubation. The trachea was cannulated with a flexible catheter, and the lungs were gently lavaged three times with 1 mL of ice-cold 0.5% bovine serum albumin (BSA) in saline; the recovered lavage fractions were pooled for cytological analysis. A total of 20,000 BALF-derived cells were cytocentrifuged onto glass slides, air-dried, and stained with hematoxylin and eosin (H&E). Slides were digitized using a Hamamatsu NanoZoomer digital slide scanner (Hamamatsu Photonics Inc., Japan) at 40× magnification (0.23 *µm*/pixel, 40× mode; equivalent to 40, 000 × 30, 000 pixels per field of view) to ensure uniform high-resolution imaging for downstream analysis. From the resulting whole-slide images (WSI), each WSI was cropped into 1122 image patches of 1024 × 1024 pixels. Patches containing *>* 80% white pixels were automatically removed while others were retained for further examination. The selection of high-quality regions followed a set of predefined criteria, including sharp cellular boundaries, minimal overlap, and well-preserved morphology. This process yielded a total of 333 high-quality image tiles, capturing a diverse representation of BALF cytology across experimental conditions (Figure 1).

**Figure 1.**
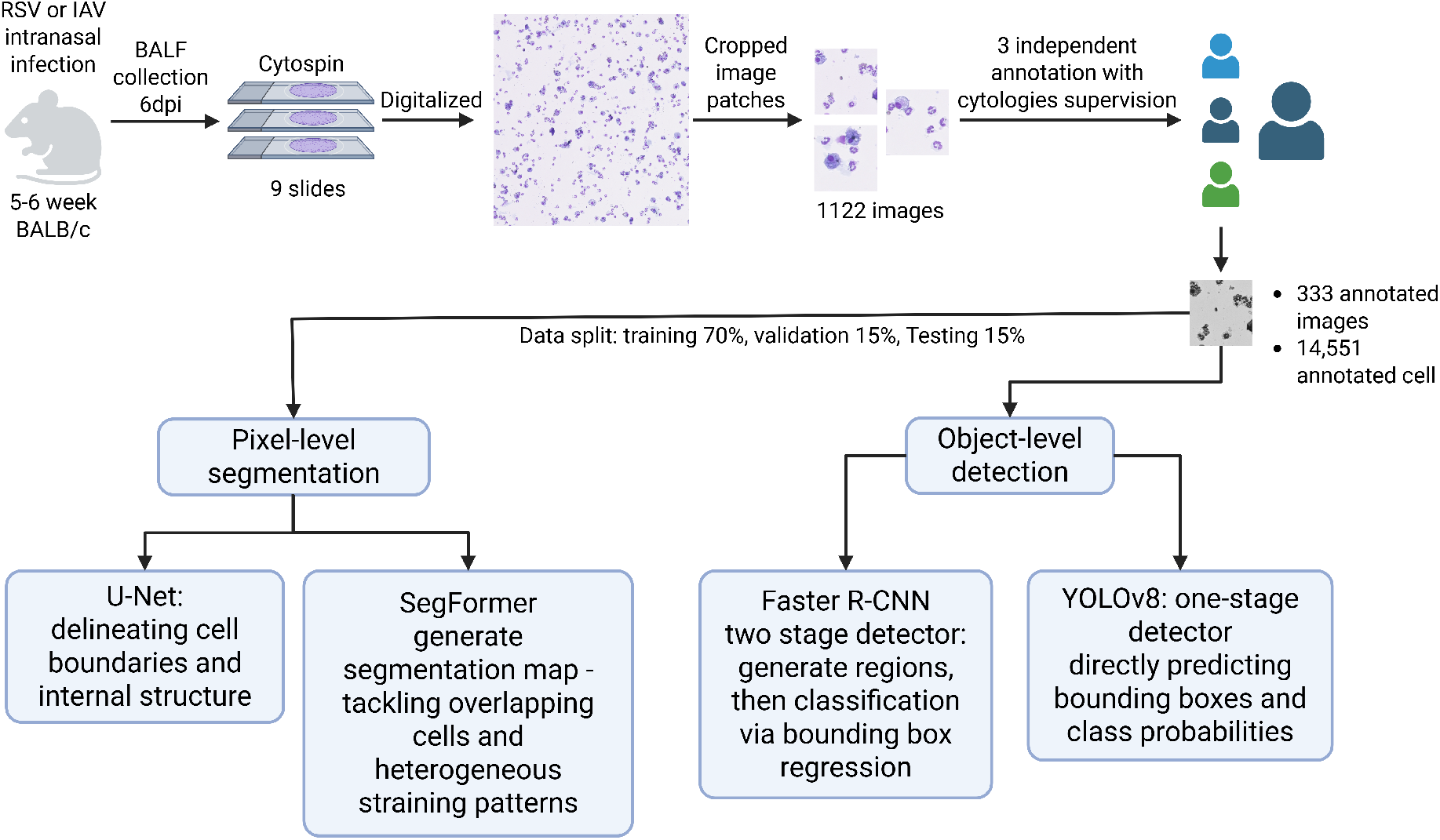
Workflow of MurineCyto-Det dataset construction and annotation. BALF samples were collected from RSV- or IAV-infected 5–6-week-old BALB/c mice at 6 days post-infection, prepared by cytospin, digitized, and cropped into image patches. After expert-supervised annotation by three independent annotators, 333 images containing 14,551 annotated cell instances were retained. The dataset was split into training, validation, and test subsets and used for two case study tasks: pixel-level segmentation with U-Net and SegFormer, and object-level detection with Faster R-CNN and YOLOv8.

### 3.2 Annotation protocol

All 333 images were annotated on the Labelbox platform using its raster-segmentation tool for pixel-level cell delineation. Five classes were defined covering the major murine BALF leukocyte populations and ambiguous material: macrophage/monocyte, neutrophil, eosinophil, lymphocyte, and unknown cell/debris. Morphological criteria for each class were:

- *Macrophage/Monocyte*: largest leukocytes; irregular or variable shape; abundant grayish-blue cytoplasm, often vacuolated; eccentric nucleus, kidney- or horseshoe-shaped.
- *Neutrophil*: multi-lobed nucleus (3–5 lobes joined by thin chromatin strands); light pink or neutral cytoplasm with fine, barely visible granules.
- *Eosinophil*: cytoplasm filled with large, uniform eosinophilic (red) granules; bilobed nucleus connected by a short chromatin bridge (spectacle-like appearance).
- *Lymphocyte*: round to slightly indented nucleus occupying most of the cell; dense chromatin; narrow rim of pale blue cytoplasm, typically agranular.
- *Unknown cell/Debris*: damaged or degenerating cells, platelet aggregates, foreign particles, or staining artifacts.

Three annotators, each trained by cytology experts, independently labeled all images. Disagreements in cell boundaries or class assignments were flagged for expert review; cytology experts adjudicated discrepancies, and this iterative review–correction cycle continued until all annotations received expert-panel approval.

Bounding boxes were derived automatically from segmentation masks using a custom script (masks2bboxes.py) that applies OpenCV contour detection (cv2.findContours) to compute the minimal bounding rectangle per cell. Masks and boxes were exported in COCO format, preserving one-to-one correspondence between segmentation and detection annotations.

### 3.3 Dataset Statistics

The MurineCyto-Det dataset comprises 333 high-resolution images of size 1024 × 1024 pixels. Across these images, a total of 14,551 cells were annotated and assigned to one of five classes: macrophage/monocyte, neutrophil, eosinophil, lymphocyte,and unknown cell/debris. Both pixel-level segmentation masks and corresponding bounding boxes are provided, yielding dual annotations for each individual cell instance.

The distribution of different cell classes is summarized in Table 1. Macrophages/monocytes (39.35%) and neutrophils (28.5%) account for the majority of annotated cells. Unknown cell/Debris (29.07%) also represents a substantial portion, reflecting the presence of damaged or unidentifiable cells and staining artifacts. In contrast, eosinophils (2.67%) and lymphocytes (0.35%) are underrepresented, creating a pronounced class imbalance within the dataset. This class imbalance represents realistic scenarios of murine BALF images, where macrophages/-monocytes and neutrophils constitute the majority of the cell population, while eosinophils and lymphocytes are considered as rare cell classes.

**Table 1.**
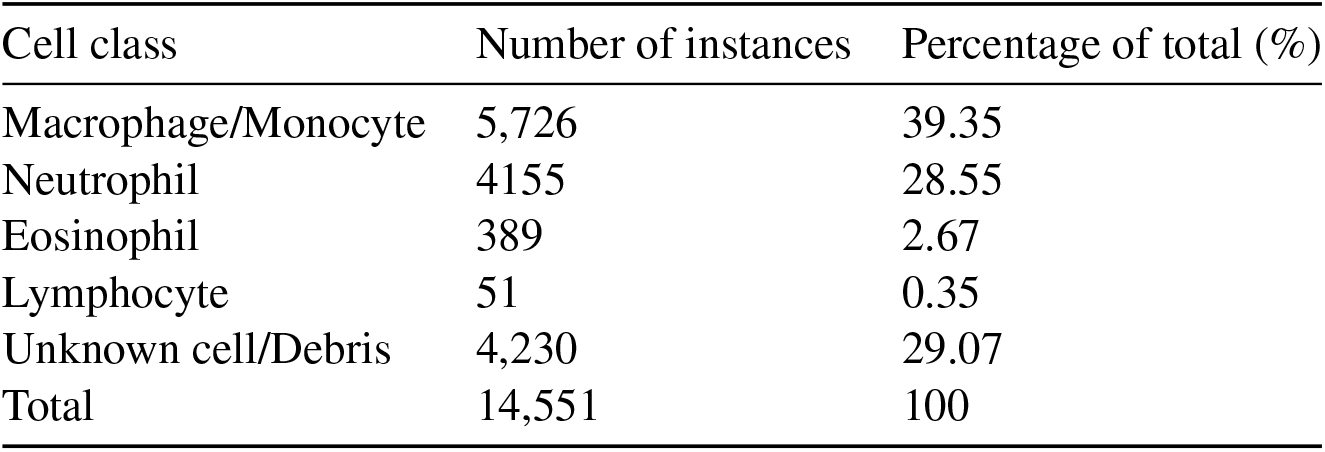
Distribution of cell classes in the dataset.

| Cell class | Number of instances | Percentage of total (%) |
| --- | --- | --- |
| Macrophage/Monocyte | 5,726 | 39.35 |
| Neutrophil | 4155 | 28.55 |
| Eosinophil | 389 | 2.67 |
| Lymphocyte | 51 | 0.35 |
| Unknown cell/Debris | 4,230 | 29.07 |
| Total | 14,551 | 100 |

In addition, the dataset exhibits considerable diversity in imaging conditions and sample characteristics. Images contain varying cell densities, from sparsely populated regions to highly crowded fields, and include cells with overlapping boundaries as well as staining artifacts. These features were deliberately retained to ensure that the dataset captures the heterogeneity encountered in realistic laboratory settings and to encourage the development of models with robust generalization capability. As shown in Table 1, this class imbalance not only reflects the biological composition of murine BALF cellularity but also poses challenges for algorithm development, as rare classes such as pulmonary infiltrating lymphocytes and eosinophils may be overlooked by models biased toward majority categories. Addressing this imbalance is therefore an important direction for future methodological innovation, for example through data augmentation, loss re-weighting, or semi-supervised learning methods (Figure 1).

Representative examples of annotated images are shown in Figure 2, illustrating the pixel-level segmentation masks and matched bounding boxes. These examples highlight the annotation quality and the diversity of cell types included in the dataset.

**Figure 2.**
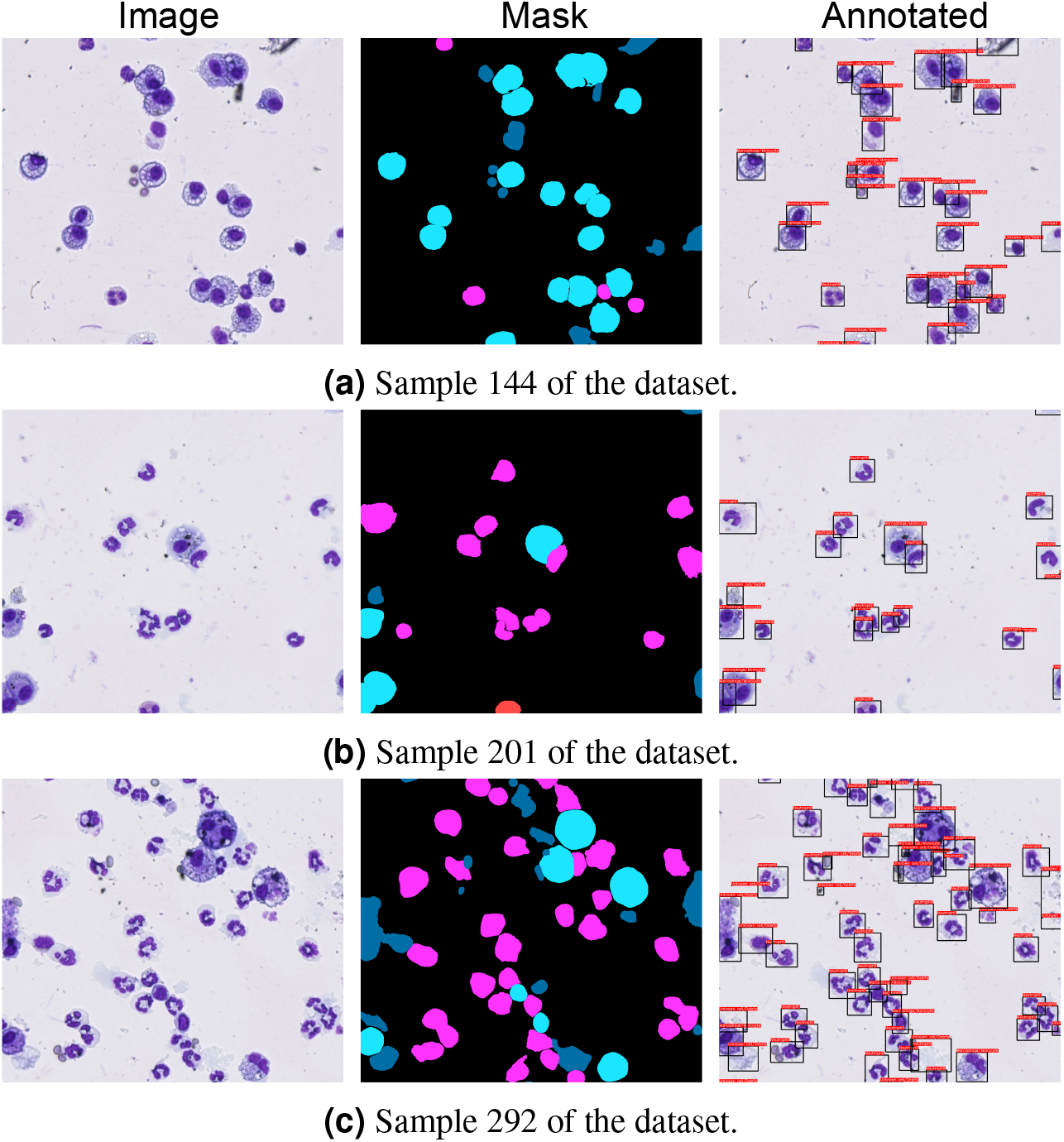
Example data samples from the dataset.

## 4 Case study

To demonstrate the practical utility of MurineCyto-Det for automated BALF cytology analysis, we established baseline benchmarks for two complementary computer vision tasks: pixel-level segmentation and object-level detection. Segmentation provides fine-grained delineation of cell extent, boundaries, and local morphology, making it suitable for downstream analyses such as morphometry and phenotype characterization. Detection instead focuses on localizing and classifying individual cells with bounding boxes and is therefore better suited to efficient cell counting and large-scale screening. Because these tasks capture the main tradeoff between morphological detail and computational efficiency, they provide a representative benchmark setting for future studies. The purpose of this case study was to establish reproducible reference baselines rather than to exhaustively compare all available architectures. Accordingly, we selected representative models that span common design choices in biomedical image analysis and evaluated all methods under a standardized experimental protocol.

### 4.1 Benchmark setup

The 333 images in MurineCyto-Det were divided into fixed training, validation, and test subsets comprising 233 images (≈ 70%), 50 images (≈ 15%), and 50 images (≈ 15%), respectively. The split was performed at the image level rather than the cell-instance level to prevent data leakage, ensuring that cells from the same image never appeared in multiple subsets. The validation set was used for model selection and early stopping, whereas the test set was reserved for final performance reporting.

For segmentation, we selected U-Net^30^ and SegFormer^31^. U-Net is a classical encoder–decoder convolutional network with skip connections and remains a standard baseline for biomedical segmentation because it preserves local spatial detail while learning semantic structure. SegFormer represents a transformer-based alternative that combines multi-scale feature extraction with lightweight decoding and is well suited to images with heterogeneous appearance and overlapping cellular structures. For detection, we selected YOLOv8^32^ and Faster R-CNN^33^. YOLOv8 is a one-stage detector that predicts bounding boxes and class probabilities in a single forward pass, making it attractive for efficient large-scale analysis. Faster R-CNN is a two-stage detector that first proposes candidate regions and then refines them, providing a strong and widely used reference framework for comparison. Together, these four models enable comparison across convolution-based versus transformer-based segmentation and one-stage versus two-stage detection paradigms.

All experiments were implemented in PyTorch version 2.7.1 with CUDA 12.9 on a workstation equipped with an NVIDIA Ada 4000 GPU (16 GB VRAM), an Intel 13900HX CPU, and 128 GB RAM under a 64-bit Windows operating system.

### 4.2 Segmentation benchmark results

For pixel-level segmentation, U-Net and SegFormer were trained using Dice loss^34^, which directly optimizes overlap between predictions and ground-truth masks. Both models used ImageNet-pretrained backbones. Optimization was performed with Adam^35^ using an initial learning rate of 1 ×10^−4^, batch size 8, and a maximum of 100 epochs. A ReduceLROnPlateau scheduler implemented in PyTorch was used to adjust the learning rate according to validation performance, and early stopping with patience 10 was applied to reduce overfitting. To improve robustness to biological and technical variability, we used a consistent augmentation pipeline comprising random horizontal flipping, affine transformations, and standardized cropping. This augmentation strategy was intended to reflect staining variation, illumination differences, and cell-shape heterogeneity commonly observed in murine cytology images.

Segmentation performance was evaluated on the test set using Pixel Accuracy (PA), mean Intersection over Union (mIoU), Dice Score^36^, and Generalized Dice Score (GDS)^37^. Formal definitions are provided in Supplementary Note 1. Class-level behavior was further examined using the confusion matrices shown in Figures 3a and 3b. Both models performed strongly on the dominant leukocyte categories, particularly macrophage/monocyte and neutrophil, indicating that MurineCyto-Det supports reliable segmentation of large and morphologically distinctive cells. U-Net achieved strong recognition of macrophage/monocyte (96%) and neutrophil (89%), with most remaining errors arising from partially overlapping regions or morphologically similar structures. Performance decreased for eosinophil, lymphocyte, and unknown cell/debris, which reflects the greater difficulty of minority and visually ambiguous classes. SegFormer showed a similar overall pattern but provided better differentiation of several challenging categories. In particular, it improved neutrophil recognition (99%) and unknown cell/debris detection (70%) relative to U-Net, although confusion between eosinophil and neutrophil remained noticeable. Lymphocyte was the most difficult category for both models, which is consistent with its limited representation in the dataset.

**Figure 3.**
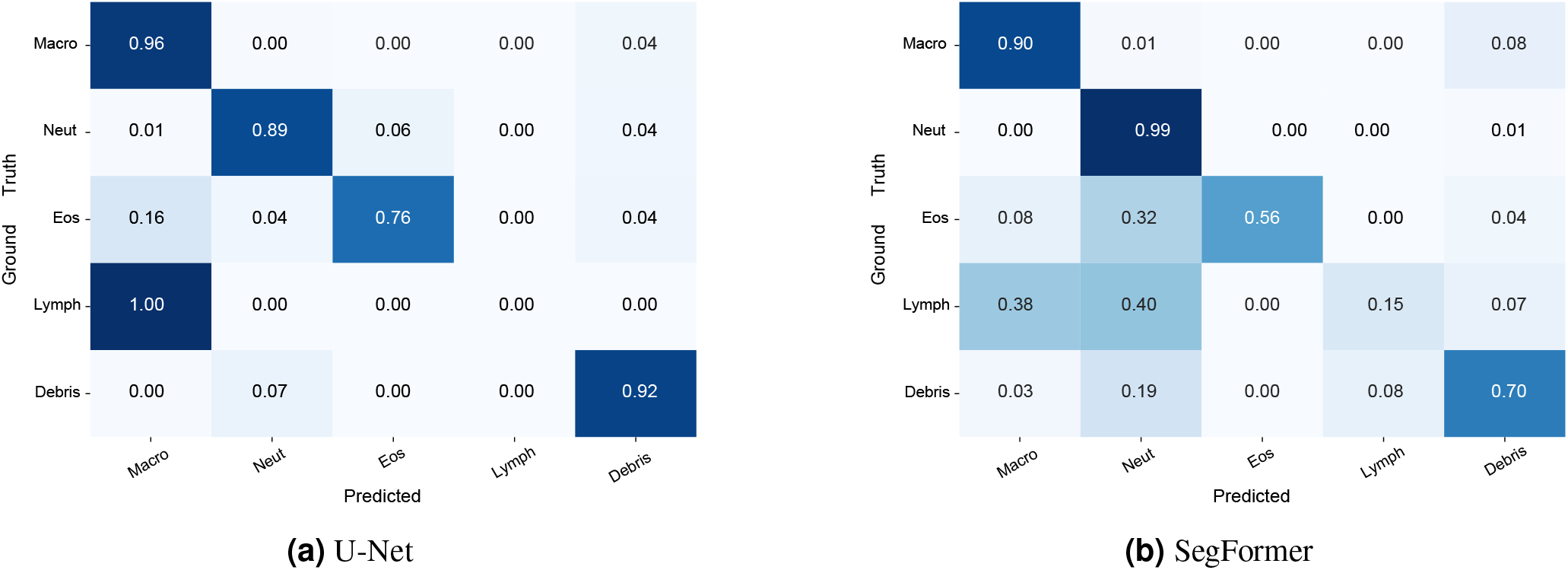
Comparison of segmentation confusion matrices on the test set. Macro - Macrophage, Neut - Neutrophils, Eos - Eosinophils, Lymph - Lymphocytes.

Representative predictions are shown in Supplementary Figure 1 and are consistent with the quantitative findings. U-Net tended to produce smooth and coherent masks for large, clearly defined cells, but it could round boundaries or merge adjacent regions when cytoplasmic areas overlapped. SegFormer more often preserved fine boundaries and structurally complex regions, which is consistent with its stronger overlap-based behavior on difficult cases. Overall, these findings show that MurineCyto-Det supports both conventional and transformer-based segmentation workflows while exposing realistic challenges associated with class imbalance and morphological heterogeneity.

### 4.3 Detection benchmark results

For object-level detection, bounding boxes were derived automatically from the segmentation masks using the one-to-one correspondence protocol described in Section 3.2. YOLOv8^32^ and Faster R-CNN^33^ were trained using the same standardized data split described above. YOLOv8 was trained with the Ultralytics PyTorch implementation using pretrained weights. Images were resized from 1024×1024 to 640×640 pixels, and training was run for 100 epochs with batch size 16 using AdamW and an initial learning rate of 1 ×10^−4^. Early stopping with patience 10 was applied based on validation performance. Faster R-CNN used a ResNet50-FPN backbone initialized with ImageNet weights and was implemented via ‘torchvision.models.detection’. Owing to the larger memory footprint of the two-stage framework, training was performed with batch size 2 using AdamW at the same initial learning rate of 1 *×* 10^−4^.

Detection performance was evaluated on the test set using precision, recall, macro-F1, mean Average Precision at IoU 0.50 (mAP@0.50), mean Average Precision at IoU 0.75 (mAP@0.75), and COCO mean Average Precision across IoU thresholds from 0.50 to 0.95 (mAP@0.50:0.95). Detailed definitions are provided in Supplementary Note 1. YOLOv8 achieved substantially stronger overall results than Faster R-CNN, with precision 0.841, recall 0.751, macro-F1 0.793, mAP@0.50 of 0.830, mAP@0.75 of 0.749, and mAP@0.50:0.95 of 0.698. The corresponding Faster R-CNN scores were 0.440, 0.439, 0.433, 0.339, 0.241, and 0.218. These results indicate that MurineCyto-Det can clearly differentiate one-stage and two-stage detector behavior under the small-object, low-contrast, and class-imbalanced conditions typical of BALF cytology images.

The class-level results in Tables 2 and 3 show the same overall trend. YOLOv8 detected macrophage/monocyte reliably and maintained moderate performance on lymphocyte and unknown cell/debris, whereas Faster R-CNN showed weaker class coverage overall and failed entirely on lymphocyte. This pattern suggests that YOLOv8 is better matched to the small, shape-variable, and morphologically subtle objects present in the dataset, whereas Faster R-CNN appears to have greater difficulty generating and refining reliable proposals for rare targets. The IoU-stratified results further support this interpretation: YOLOv8 achieved mAP@0.75 of 0.749, compared with 0.241 for Faster R-CNN, indicating substantially stronger localization quality once detections were made. Although both models showed reduced performance at stricter IoU thresholds, YOLOv8 remained consistently more accurate across the full evaluation range.

**Table 2.** Per-class metrics of YOLOv8.

| Class | Prec. | Rec. | F1 |
| --- | --- | --- | --- |
| Macro/Mono | 0.928 | 0.934 | 0.931 |
| Eosinophil | 0.91 | 0.854 | 0.881 |
| Neutrophil | 0.94 | 0.758 | 0.839 |
| Lymphocyte | 0.75 | 0.545 | 0.631 |
| Unknown | 0.679 | 0.664 | 0.671 |

**Table 3.** Per-class metrics of Faster R-CNN.

| Class | Prec. | Rec. | F1 |
| --- | --- | --- | --- |
| Macro/Mono | 0.352 | 0.418 | 0.382 |
| Neutrophil | 0.841 | 0.593 | 0.696 |
| Eosinophil | 0.543 | 0.667 | 0.598 |
| Lymphocyte | 0 | 0 | 0 |
| Unknown | 0.467 | 0.518 | 0.491 |

Taken together, these results provide practical and reproducible benchmark baselines for MurineCyto-Det and standardized reference points for the development, evaluation, and comparison of future methods on this dataset.

## 5 Conclusion

We introduced MurineCyto-Det, a high-resolution murine BALF cytology dataset designed to support reproducible automated cell analysis in preclinical respiratory research. The dataset contains 333 image tiles with 14,551 annotated cell instances across five cytological categories, together with expert-validated pixel-level segmentation masks and one-to-one matched bounding boxes. By providing paired segmentation and detection annotations, MurineCyto-Det offers a standardized resource for evaluating automated cytology methods under consistent experimental conditions. The benchmark results demonstrate that MurineCyto-Det supports both morphology-oriented segmentation and efficient object-level detection. U-Net and SegFormer provided representative segmentation baselines, showing that the dataset can support pixel-level analysis while also revealing challenges associated with overlapping cells, ambiguous boundaries, and class imbalance. YOLOv8 and Faster R-CNN provided complementary detection baselines, with YOLOv8 achieving stronger overall detection and localization performance under the small-object and morphologically variable conditions present in murine BALF cytology images. These findings indicate that MurineCyto-Det is sufficiently informative for practical model evaluation while remaining challenging enough to expose meaningful differences among computational approaches. Overall, MurineCyto-Det fills an important resource gap for automated murine BALF cytology by combining expert annotations, dual task support, and reproducible benchmark baselines. The dataset and accompanying code are publicly released to facilitate future method evaluation and comparison. Future work may build on this resource through data-efficient learning, semisupervised annotation strategies, domain adaptation between murine and human cytology, and integration with multimodal imaging or experimental metadata.

## Author contributions

L.D.V.and Y.D. conceptualized and directed the study. T.X.L., D.A.F., S.W., and Y.D. developed and implemented the coding pipeline. L.D.V., L.-A.T.T., A.T.Q.P., and S.A.C. performed mouse experiments and analyzed data. A.T.L.T., L.D.V., A.T., and D.M. performed data labeling. T.X.L., Y.D., L.-A.T. T. and L.D.V. drafted the manuscript.

## Data availability

The MurineCyto-Det dataset is publicly available through Zenodo at at https://doi.org/10.5281/zenodo.17608677 under the Creative Commons Attribution 4.0 International License (CCBY 4.0). The case study code, training scripts, split files, and evaluation scripts are available at the accompanying GitHub repository https://github.com/Le-Xuan-Thang/MurineCyto-Det.

