## Supplementary Note 1 and Supplementary Figure 1 for "MurineCyto-Det: A High-Resolution Murine BALF Cytology Dataset for Leukocyte Segmentation and Detection"

#### Table of contents

**Supplementary Notes** .......... 2

**Supplementary Figures** .......... 4

### Supplementary Notes

#### Supplementary Note 1: Evaluation Metrics

For the segmentation task, performance was evaluated using four widely adopted metrics: Pixel Accuracy (PA), Mean Intersection over Union (mIoU), Dice Score, and Generalized Dice Score (GDS). In addition, the Dice loss was employed as the optimization objective during training.

Pixel Accuracy (PA) measures the proportion of correctly classified pixels over all pixels in the dataset. It provides a simple overall measure of performance but may be biased toward majority classes when class imbalance is present. Formally:

$$PA = \frac{\sum_i n_{ii}}{\sum_i t_i} \tag{1}$$

where $n_{ii}$ is the number of correctly predicted pixels for class $i$, and $t_i$ is the total number of pixels belonging to class $i$.

The Intersection over Union (IoU) quantifies the overlap between predicted and ground-truth regions for each class. The mean IoU averages this score across all $C$ classes:

$$mIoU = \frac{1}{C} \sum_{i=1}^{C} \frac{n_{ii}}{t_i + \sum_j n_{ji} - n_{ii}} \tag{2}$$

where $n_{ii}$ is the number of correctly predicted pixels of class $i$, $t_i$ is the number of ground-truth pixels of class $i$, and $n_{ii}$ represents the number of pixels predicted as class $i$ but belonging to class $j$.

The Dice Score coefficient (equivalent to the F1-score in binary segmentation) measures the similarity between predicted and ground-truth masks. For class $i$, it is defined as:

$$\text{Dice}_{\ i} = \frac{2\,|P_i \cap G_i|}{|P_i| + |G_i|} \tag{3}$$

where $P_i$ and $G_i$ denote the predicted and ground-truth pixel sets for class $i$. The average Dice score across classes is reported.

The Generalized Dice Score (GDS) extends the Dice coefficient to mitigate the effect of class imbalance by weighting each class according to the inverse of its frequency. It is computed as:

$$GDS = \frac{2\sum_{i=1}^{C} w_i\,|P_i \cap G_i|}{\sum_{i=1}^{C} w_i\,(|P_i| + |G_i|)} \tag{4}$$

where $w_i = \frac{1}{(|G_i|)^2}$ is the weight assigned to class $i$, inversely proportional to the square of its ground-truth pixel count. This weighting scheme increases the contribution of rare classes to the overall score.

For the object detection task, model performance was evaluated using standard metrics derived from the Precision–Recall (PR) curve and Intersection over Union (IoU) overlap between predicted and ground-truth bounding boxes. Following the COCO and PASCAL VOC conventions, the main quantitative indicators include mean Average Precision (mAP), Precision, Recall, and $F_1$-score. These metrics jointly capture the trade-off between localization quality and classification reliability.

For a predicted bounding box $B_p$ and its corresponding ground-truth box $B_g$, the IoU is defined as:

$$\text{IoU} = \frac{|B_p \cap B_g|}{|B_p \cup B_g|}, \tag{5}$$

where $|\cdot|$ denotes the area in pixels. A prediction is considered correct if IoU $\geq \tau$, with $\tau$ typically set to 0.5 (i.e., IoU@50).

Precision ($P$) and Recall ($R$) are defined as:

$$P = \frac{TP}{TP + FP}, \qquad R = \frac{TP}{TP + FN}, \tag{6}$$

where $TP$, $FP$, and $FN$ represent the number of true positives, false positives, and false negatives, respectively. Precision measures the proportion of correct detections among all predicted boxes, while Recall measures the proportion of correctly detected objects among all ground-truth instances.

The Average Precision (AP) for a given class is computed as the area under the Precision–Recall (PR) curve:

$$AP = \int_0^1 P(R)\, dR, \tag{7}$$

where $P(R)$ denotes precision as a function of recall. The mean Average Precision (mAP) aggregates the $AP$ across all classes $C$:

$$\text{mAP} = \frac{1}{C} \sum_{i=1}^{C} AP_i. \tag{8}$$

To evaluate localization robustness, three thresholds were used:

- mAP@50: average precision at IoU $\geq 0.50$, representing coarse localization accuracy.
- mAP@75: average precision at IoU $\geq 0.75$, emphasizing stricter bounding-box alignment.
- mAP@[50 : 95]: COCO-style mean average precision averaged over ten IoU thresholds (0.50:0.05:0.95), providing a comprehensive measure of performance across localization tolerances.

The harmonic mean of precision and recall provides a single scalar measure of detection quality:

$$F_1 = 2 \times \frac{P \times R}{P + R}. \tag{9}$$

To ensure class balance, a macro-averaged $F_1$-score was computed across all $C$ classes:

$$F_{1,macro} = \frac{1}{C} \sum_{i=1}^{C} F_{1,i}. \tag{10}$$

This metric treats all classes equally, mitigating the influence of dominant categories and highlighting rare cell types such as lymphocyte and eosinophil.

### Supplementary Figures

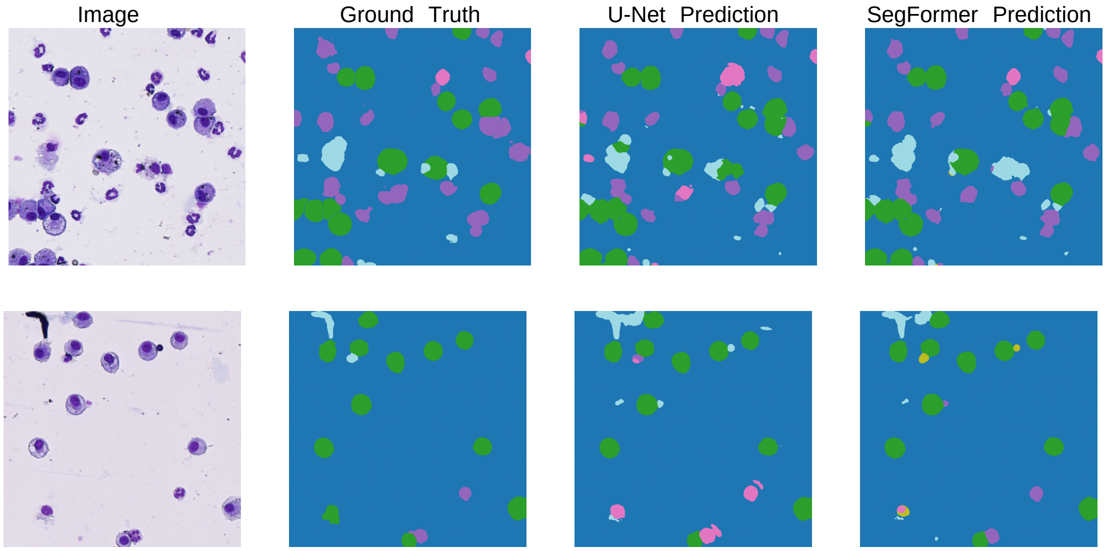


**Supplementary Figure 1.** Examples of prediction results from the two models.
